# Genome-wide CNV study and functional evaluation identified *CTDSPL* as tumour suppressor gene for cervical cancer

**DOI:** 10.1101/353391

**Authors:** Dan Chen, Shuai Wang, Zhenli Li, Tao Cui, Yao Chen, Junjiao Song, Patrik KE Magnusson, Huibo Wang, Dandan Zhang, Ulf Gyllensten

**Author notes:** These authors contribute equally to the work (Dan Chen, Shuai Wang, Zhenli Li, Tao Cui). Correspondence. Correspondence and material requests should be addressed to 1. Dan Chen. Ministry of Education and Shanghai Key Laboratory of Children’s Environmental Health, Xinhua Hospital, Shanghai Jiao Tong University School of Medicine, 1665 Kongjiang Road, 200092 Shanghai, China.; 2.Huibo Wang. Department of Neurosurgery, First Affiliated Hospital of Nanjing Medical University, Nanjing, Jiangsu, China.; 3. Dandan Zhang. Department of Pathology, School of Medicine, Zhejiang University, Hangzhou, Zhejiang, China.

## Abstract

We have investigated copy number variations (CNVs) in relation to cervical cancer by analyzing 731,422 single-nucleotide polymorphisms (SNPs) in 1,034 cervical cancer cases and 3,948 controls, followed by replication in 1,396 cases and 1,057 controls. We found that a 6367bp deletion in intron 1 of the CTD small phosphatase like gene (*CTDSPL*) was associated with 2.54-fold increased risk of cervical cancer (odds ratio =2.54, 95% confidence interval =2.08-3.12, *P*=2.0×10^−19^). This CNV is one of the strongest genetic risk variants identified so far for cervical cancer. The deletion removes the binding sites of zinc finger protein 263, **binding protein 2 and interferon regulatory factor 1**, and hence downregulates the transcription of *CTDSPL*. HeLa cells expressing *CTDSPL* showed a significant decrease in colony-forming ability. Compared with control groups, mice injected with HeLa cells expressing *CTDSPL* exhibited a significant reduction in tumour volume. Furthermore, *CTDSPL*-depleted immortalized End1/E6E7 could form tumours in NOD-SCID mice.

## Introduction

Cervical cancer is the third most common cancer in women worldwide (Arbyn et al. 2011). Genome-wide association studies (GWASs) focusing on single-nucleotide polymorphisms (SNPs) have identified multiple genetic susceptibility loci for cervical cancer (Chen et al. 2013; Shi et al. 2013). However, the risk variants identified to date have small effect sizes (per allele odds ratio (OR) < 1.50) and only explain a small fraction of the heritability. Although epistatic and gene-environment interactions may contribute to the unexplained heritability of cervical cancer, it seems likely that a significant fraction is due to loci that have not yet been identified. Recent studies indicate that copy number variations (CNVs) occur frequently in the genome and are an important source of human genetic variation (Iafrate et al. 2004; Sebat et al. 2004). It has been proposed that CNVs may explain some of the missing heritability for complex diseases after the findings from GWASs (Eichler et al. 2010).

The role of somatic CNVs in cervical cancer has been extensively studied (Snijders et al. 1998; Ng et al. 2007; Mitra et al. 2010a; Lee et al. 2012; Mendoza-Rodriguez et al. 2013; Lee et al. 2014; Zhou et al. 2017). However, very few studies have evaluated the association of germline CNVs with cervical cancer risk. Only one study demonstrated that a lower defensin beta 104A (*DEFB4*) copy number was associated with susceptibility to cervical cancer (Abe et al. 2013). To assess the association of both common and rare germline CNVs with cervical cancer risk, we conducted a genome-wide CNV study of 1,034 cervical cases and 3,948 controls in a Swedish population using Illumina HumanOmniExpress BeadChip (Illumina, San Diego, CA) (731,422 SNPs) followed by a replication study of 1,396 cervical cancer subjects and 1,057 controls.

## Results

### Association between CNVs and cervical cancer risk

Principal component analysis (PCA) using 17,386 markers in low linkage disequilibrium (LD) (pair-wise *r*^2^<0.02) showed that there was no statistically significant difference between cases and controls (*P*=0.330) (**Figure 1**), suggesting no batch effect. Meanwhile, the quantile-quantile plot shows minimal evidence of genomic inflation (*λ*.=1.031), suggesting no systematic bias (Chen et al. 2013). To maximize the finding of potential cervical cancer-associated CNVs, the two algorithms-QuantiSNP 2.0 (Colella et al. 2007) and PennCNV (Wang et al. 2007) were used for identifying CNVs from the signal intensity data of the SNP microarray (log-R ratio and B-allele frequency). After initial quality control (**Methods**), 973 cases with 11,056 autosomal CNVs and 3,485 controls with 33,492 autosomal CNVs were included for global burden analysis in the discovery stage. In order to improve prediction accuracy, as to every specific gene/region that we were interested, samples with inconsistent CNV calls from two algorithms were further removed. Therefore, the number of samples included in the final analysis varied across specific genes/regions.

**Figure 1.**
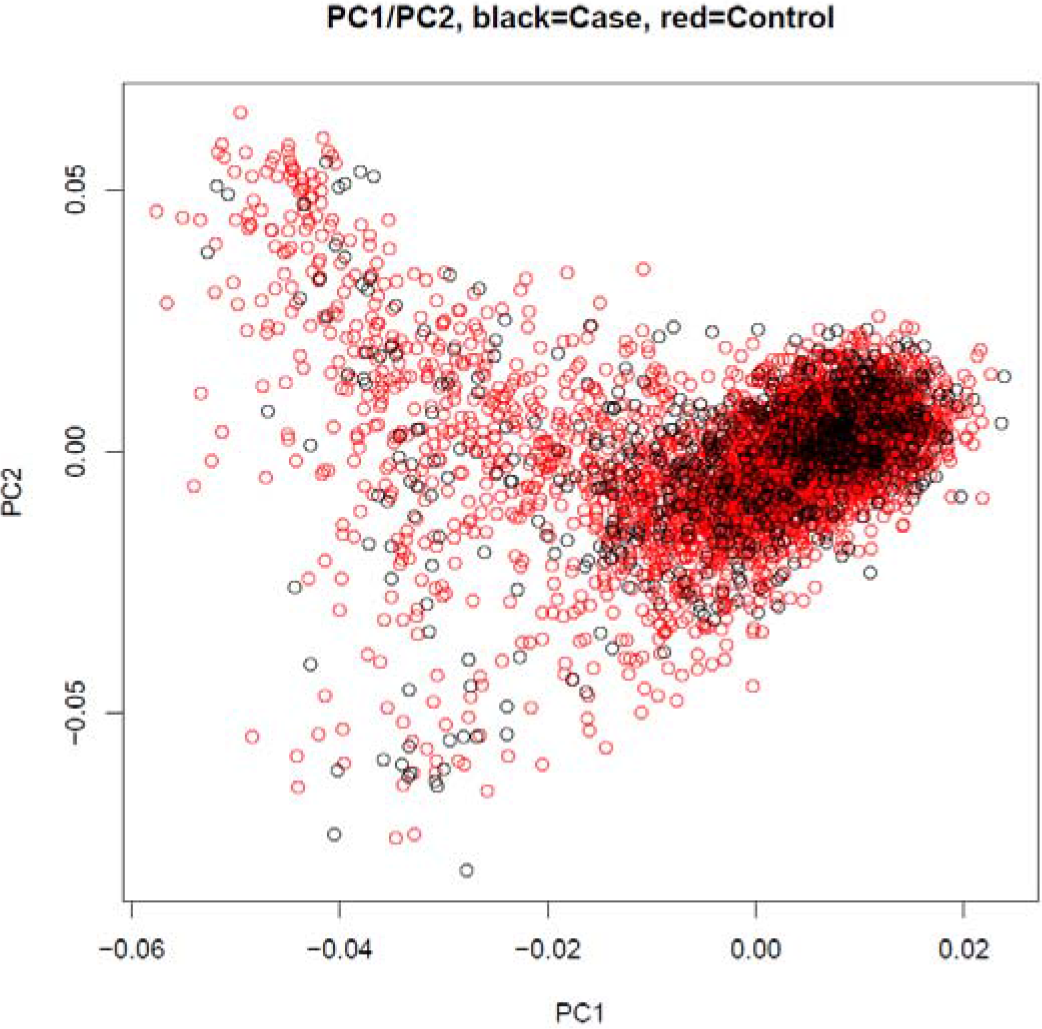
Principal component (PC)1 plotted against PC2 generated by principal component analysis (PCA). The black circles represent cervical cancer patients and the red circles represent controls.

Cervical cancer patients had a higher total genome-wide burden of CNVs (all categories), all genic CNVs, rare CNVs and rare genic CNVs than controls (fold change: 1.18, 1.16, 1.19, 1.09 respectively, all *P*< 0.05, **Table 1**). The difference between patients and controls was particularly strong for duplications (fold change: 1.24, 1.22, 1.25, 1.19 for all duplications, all genic duplications, rare duplications and rare genic duplications respectively, all with a *P*<0.05, **Table 1**). All analyses, except for rare genic deletions (fold change=1.05, *P* = 0.13), reached statistical significance (*P* < 0.05).

**Table 1.** Frequency of CNVs in cases versus controls.

| Type | Category | Cases (%) | Control (%) | Fold change | *P* value |
| --- | --- | --- | --- | --- | --- |
| Total | All | 11.36 | 9.64 | 1.18 | $<1.0 \times 10^{-6}$ |
| | All genic | 4.28 | 3.69 | 1.16 | $<1.0 \times 10^{-6}$ |
| | Rare | 3.29 | 2.76 | 1.19 | $<1.0 \times 10^{-6}$ |
| | Rare genic | 1.45 | 1.33 | 1.09 | $4.4 \times 10^{-3}$ |
| Deletions | All | 9.15 | 7.85 | 1.17 | $<1.0 \times 10^{-6}$ |
| | All genic | 3.13 | 2.74 | 1.14 | $<1.0 \times 10^{-6}$ |
| | Rare | 2.35 | 2.00 | 1.17 | $<1.0 \times 10^{-6}$ |
|  | Rare genic | 0.97 | 0.92 | 1.05 | 0.13 |
| Duplications | All | 2.21 | 1.79 | 1.24 | $<1.0 \times 10^{-6}$ |
| | All genic | 1.15 | 0.95 | 1.22 | $<1.0 \times 10^{-6}$ |
| | Rare | 0.94 | 0.75 | 1.25 | $<1.0 \times 10^{-6}$ |
| | Rare genic | 0.48 | 0.41 | 1.19 | $8.0 \times 10^{-4}$ |

*P* values were inferred using $1 \times 10^6$ permutations by PLINK and statistically significant *P* values ( $P < 0.05$ ) were shown in bold.

We then explored whether individual genes impacted by deletions or duplications were associated with susceptibility to cervical cancer. The strongest association was found for a 6367bp deletion in intron 1 of the CTD small phosphatase like gene (*CTDSPL*) (chr3: 37979882- 37986249), which was found in 7.4 % of cervical cancer cases and 2.6 % of controls (odds ratio (OR)=2.61, 95% confidence interval (CI)=1.91-3.56, *P*=1.5×10^−9^, BFDP <0.5 when the prior probability is 0.5 or 0.05) (**Table 1**). Associations with deletions in *ATP13A4* (OR=2.31, 95% CI=1.55-3.45, *P*= 4.3×10^−5^) and *NEDD4L* (OR=4.09, 95% CI =2.07-8.05, *P*= 4.6×10^−5^) also reached statistical significance but failed correction for multiple testing when the prior probability is 0.05 and 0.5, respectively (**Table 2**). To verify the accuracy of CNV calls, we reanalyzed a proportion of samples using both real-time quantitative PCR (RT-qPCR) and Custom Taqman Copy Number Assays (Life Technologies). Deletion of *ATP13A4* was excluded from further investigation as it was unable to be designed for RT-qPCR or Taqman assay. Both the SybrGreen RT-qPCR and Custom Taqman Assays showed 100% concordance with the CNV calls from GWAS data on 99 cervical cancer cases tested for *CTDSPL* and 75 cervical cancer cases tested for *NEDD4L* based on the SNP arrays.

**Table 2.** CNV-disrupted genes associated with cervical cancer identified in the discovery phase.

| CNV | Position* | Type | Frequency |  | EMP<sup>†</sup> | Association<sup>‡</sup> |  |  | BFDP<sup>§</sup> |  |
| --- | --- | --- | --- | --- | --- | --- | --- | --- | --- | --- |
|  |  |  | Case | Control |  | OR | 95%CI | P | 0.5 | 0.05 |
| ***CTDSPL*** | chr3: 37979882-37986249 | deletion | 0.07 | 0.03 | $<1.0 \times 10^{-6}$ | 2.61 | 1.91-3.56 | $1.5 \times 10^{-9}$ | $3.0 \times 10^{-4}$ | 0.03 |
| ***ATP13A4*** | chr3:193119865-193272696 | deletion | 0.04 | 0.02 | $1.0 \times 10^{-5}$ | 2.31 | 1.55-3.45 | $4.3 \times 10^{-5}$ | 0.28 | 0.98 |
| ***NEDD4L*** | chr18:55711779-56065389 | deletion | 0.02 | 0.005 | $7.0 \times 10^{-5}$ | 4.09 | 2.07-8.05 | $4.6 \times 10^{-5}$ | 0.71 | 1.00 |

OR, Odds ratio; CI, confidence interval; BFDP, Bayes false discovery probability.
\* Genomic positions were based on human genome assembly 19 (hg19).
† EMP represented *P* values calculated by PLINK using permutation tests (1,000,000 iterations).
‡ Odds ratio, 95% confidence interval and *P* values for the deletions were calculated by logistic regression in the allelic test using SAS.
§ Calculated as described previously (21).

The association with deletion in *CTDSPL* was further replicated in an independent study of 1,396 cervical cancer subjects and 1,057 controls (OR=1.31, 95%CI=1.05-1.65, *P*=0.017) from the Swedish population (**Methods**, Table 3). The frequency of the deletion is 0.17 and 0.14 in the cancer patients and controls, respectively. In contrast, no association was observed between deletion in *NEDD4L* and cervical cancer risk in the replication series (OR=1.04, 95% CI=0.48-2.26, *P*=0.92). Using the combined discovery and replication data, the deletion in *CTDSPL* showed an OR = 2.54 (95% CI = 2.08-3.12, *P* = 2.0×10^−19^) with risk of cervical cancer. No heterogeneity for the association of *CTDSPL* deletion with cervical cancer risk was noted by tumour stage (CIS vs invasive cancer) (*P* = 0.12).

**Table 3.** Association results of *CTDSPL*-6367bp deletion with cervical cancer risk in the combined analysis.

|  | Number (Frequency) |  | Association* |  |  |
| --- | --- | --- | --- | --- | --- |
|  | Case | Control | OR | 95%CI | P |
| **Overall** | 2290(0.13) | 4245(0.05) | 2.54 | 2.08-3.12 | 2.0×10<sup>-19</sup> |
| **By genotype** |  |  |  |  |  |
| 0 copy of del | 1990(0.87) | 4016(0.95) | Ref | Ref |  |
| 1 copies of del | 272(0.12) | 208(0.05) | 2.64 | 2.14-3.26 | 1.6×10<sup>-19</sup> |
| 2 copies of del | 28(0.01) | 21(0.005) | 2.69 | 0.81-8.97 | 0.10 |
| **By study** |  |  |  |  |  |
| Discovery | 894(0.07) | 3188(0.03) | 2.61 | 1.91-3.56 | 1.5×10<sup>-9</sup> |
| Replication | 1396(0.17) | 1057(0.14) | 1.31 | 1.05-1.65 | 0.02 |
| **By tumour stage** |  |  |  |  |  |
| Carcinoma *in situ* | 2096(0.13) | 4245(0.05) | 2.45 | 1.99-3.02 | 3.7×10<sup>-17</sup> |
| Invasive cancer | 194(0.19) | 4245(0.05) | 3.58 | 2.32-5.53 | 8.6×10<sup>-9</sup> |

del, deletion; OR, odds ratio; CI, confidence interval
\* Odds ratio, 95% confidence interval and *P* values for the deletions were calculated by Pedgenie which corrected for relatedness except for the discovery phase in which odds ratio, 95% confidence interval and *P* values were calculated by logistic regression in the allelic test as there were no related subjects.

### The effect of *CTDSPL* deletion on transcription

RNA-seq data in cervical cancer tissues from 293 individuals in The Cancer Genome Atlas (TCGA) (http://www.cancergenome.nih.gov) indicated that the 6367bp deletion was significantly associated with decreased expression of *CTDSPL* (Spearman coefficient = 0.54, *P* = 1.4×10^−23^), supporting that the deletion affects the transcription of *CTDSPL* (**Methods**). There are 3 other genes (villin like (*VILL*), phospholipase C delta 1(*PLCD1*) and deleted in lung and esophageal cancer 1 (*DLEC1*)) within the 200kb range of the deletion. We analyzed the data from TCGA to examine the association between the *CTDSPL* CNV and the expression level of these candidate genes (Methods). One-way ANOVA results suggested that expression level of *CTDSPL* showed more significant association with CNV of *CTDSPL* than the other genes (7.3 ×10^−19^ vs 0.02, 0.001 and 0.75, respectively) (**Figure 2**, **Table 4**). In addition, the significance of the correlation with *VILL* and *PLCD1* did not remain in the linear regression analysis, while *CTDSPL* itself still showed remarkable significance (5.1×10^−9^, **Table 5**). Therefore, we speculated that the deletion of *CTDSPL* may be more likely to affect the transcription of *CTDSPL* itself rather than the surrounding genes.

**Figure 2.**
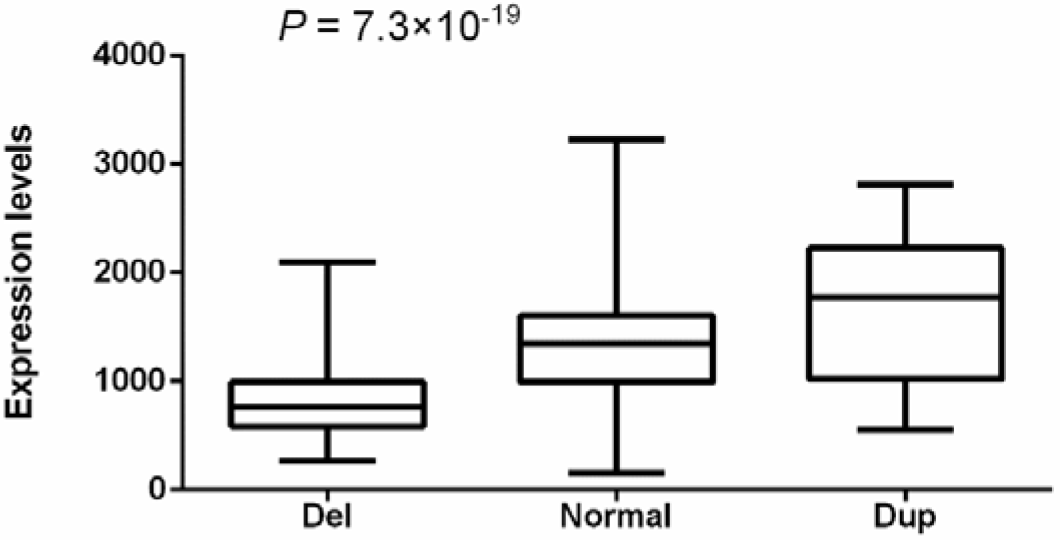
Correlation between CNV and *CTDSPL* transcriptional level. 293 cervical squamous cell carcinoma and endocervical adenocarcinoma (CESC) from The Cancer Genome Atlas (TCGA) (http://www.cancergenome.nih.gov) with both copy number variation data and RNA-seq data available were included. Statistical significance between groups was assessed using one-way ANOVA.

**Table 4.** The association between the deletions that cover the 6367bp fragment in *CTDSPL* and expression levels of surrounding genes.

| **Gene** | **Deletion (N=91)** | **Normal (N=187)** | **Duplication (N=15)** | **P** |
| --- | --- | --- | --- | --- |
| ***CTDSPL*** | 840.7 ± 379.8 | 1380.2 ± 555.8 | 1712.3 ± 653.0 | **7.3×10<sup>-19</sup>** |
| ***VILL*** | 438.6 ± 619.7 | 713.9 ± 884.0 | 683.4 ± 705.2 | **0.02** |
| ***PLCD1*** | 396.3 ± 302.1 | 512.9 ± 523.7 | 861.4 ± 868.1 | **0.001** |
| ***DLEC1*** | 40.4 ± 227.3 | 28.2 ± 105.0 | 15.5 ± 34.0 | 0.75 |

<sup>a</sup> Calculated by one-way ANOVA.

**Table 5.**
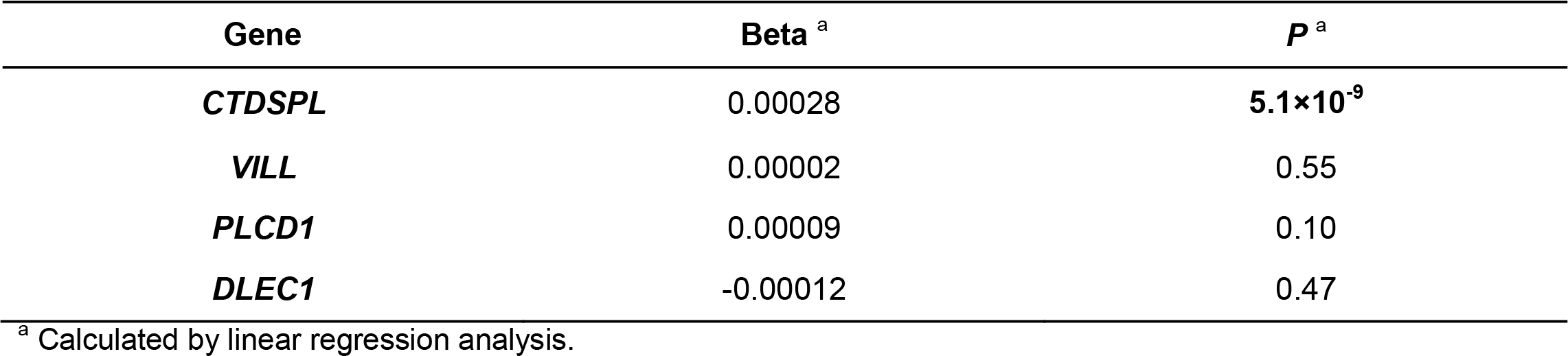
Linear regression analysis between the deletions that cover the 6367bp fragment in *CTDSPL* and expression levels of surrounding genes.

As shown in **Figure 3**, according to the Encyclopedia of DNA Elements (ENCODE) data (Consortium 2012), 6 transcriptional factors were predicted to bind to the *CTDSPL* intron1-6367bp fragment (chr3:37979882-37986249), i.e upstream transcription factor 2 (USF2) (chr3:37,981,985-37,982,248), activating transcription factor 3 (ATF3) chr3:37,982,058-37,982,290), upstream transcription factor 1 (USF1) (chr3:37,982,074-37,982,317), zinc finger protein 263 (ZNF263) (chr3:37,982,058-37,982,290); binding protein 2 (GATA2) (chr3:37,983,211-37,983,617) and interferon regulatory factor 1(IRF1) (chr3:37,986,122- 37,986,432). All of them are expressed in cervical cells (Klijn et al. 2015). Three DNase-sensitive sites were found to be overlapped with the binding sites of the above five transcriptional factors, respectively, which are usually regulatory regions in general, and promoters in particular (Consortium 2012). ZNF263, GATA2 and IRF1 have been implicated in cancer development (Willman et al. 1993; Harada et al. 1994; Tamura et al. 1996; Nozawa et al. 1998; Nozawa et al. 1999; 22021428Bohm et al. 2009; Frietze et al. 2010; Kazenwadel et al. 2012; Kumar et al. 2012; Vicente et al. 2012; Wang et al. 2012; Chiang et al. 2014; He et al. 2014; Klijn et al. 2015; Wang et al. 2015; Chen et al. 2017).

**Figure 3.**
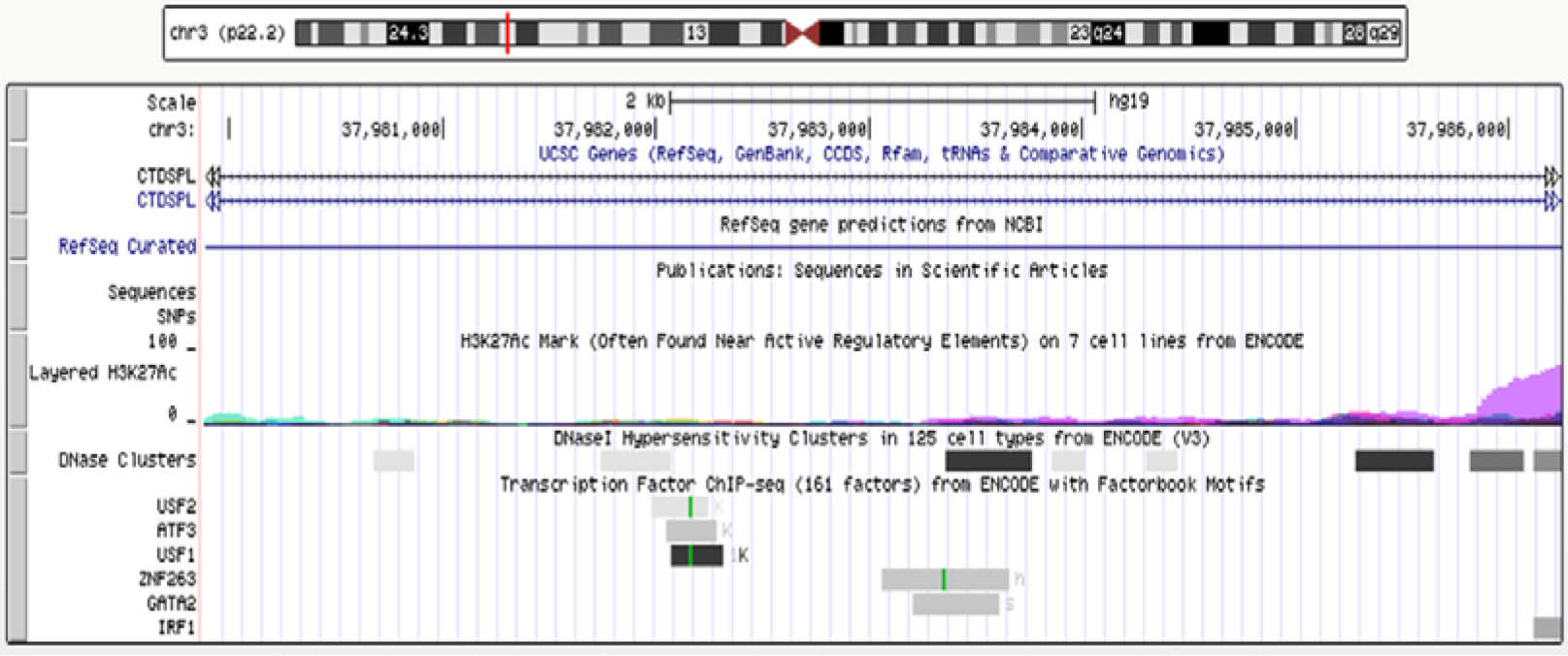
Genomic region at chr3:37979382-37986749 (http://genome.ucsc.edu/cgi-bin/hgTracks?db=hg19&lastVirtModeType=default&lastVirtModeExtraState=&virtModeTvpe=default&virtMode=0&nonVirtPosition=&position=chr3%3A37979882-37986249&hgsid=667990619b8hObVvlmaFxQft1gH4IdnRAH2Vz.) The track of “layered H3K27Ac” shows the levels of enrichment of the H3K27Ac histone mark across the genome as determined by a ChIP-seq assay. The H3K27Ac histone mark is the acetylation of lysine 27 of the H3 histone protein, and it is thought to enhance transcription possibly by blocking the spread of the repressive histone mark H3K27Me3. The track of “DNase clusters” shows DNase hypersensitive areas assayed in a large collection of cell types by the ENCODE project. Regulatory regions in general, and promoters in particular, tend to be DNase-sensitive. The track at the bottom shows regions of transcription factor binding derived from a large collection of ChIP-seq experiments performed by the ENCODE project, together with DNA binding motifs identified within these regions by the ENCODE Factorbook repository (24).

The effect of the location of this fragment on transcriptional regulation was examined by luciferase reporter assays using two constructs containing the fragment upstream and downstream of the SV40 promoter, respectively (**Methods**, **Figure 4A**). Relative promoter activity was determined in HEK293T cells that over and under-express ZNF263, respectively. Both constructs containing the *CTDSPL*-6367bp fragment upstream or downstream of the SV40 promotor generated higher luciferase signals as compared to PGL3-Controls without the *CTDSPL*-6367bp fragment, when transfecting 293T which overexpressed ZNF263 through lentiviral plasmid construction (*P* = 0.04, *P* = 0.03, respectively) (**Figure 4B**), suggesting that the location of the *CTDSPL*-6367bp fragment relative to the SV40 promoter (upstream or downstream) did not affect the transcriptional activity. However, there was no significant difference in the luciferase signal between constructs containing the fragment and PGL3-Control when transfecting 293T that did not overexpress ZNF263 (*P* > 0.05) (**Figure 4C**). Then we examined relative promoter activity using construct containing the fragment upstream of the SV40 promoter in HEK293T cells that overexpress ZNF263, GATA2 and IRF1, respectively. Flavin containing monooxygenase 3 (FMO3), a transcription factor predicted to have no binding site in intron 1 of *CTDSPL* and is expressed in cervical cells (Klijn et al. 2015), was set as an additional control. The construct containing the CTDSPL-6367bp fragment upstream of the SV40 promotor generated higher luciferase signals as compared to PGL3-Control without the *CTDSPL*-6367bp fragment, when co-transfected with *ZNF263*, *GATA2* or *IRF1* cDNA to HEK293T cells (*P* = 0.006, *P* = 0.035, *P* = 0.016, respectively) (**Figure 5**). However, there was no significant difference in the luciferase signals between the construct containing the *CTDSPL*-6367bp fragment and PGL3-Control when co-transfected with *FMO3* cDNA to HEK293T (*P* > 0.05). These results indicate that the enhancer activity of the *CTDSPL* intron 1-6367bp fragment depends on the expression of ZNF263, GATA2 or IRF1.

**Figure 4.**
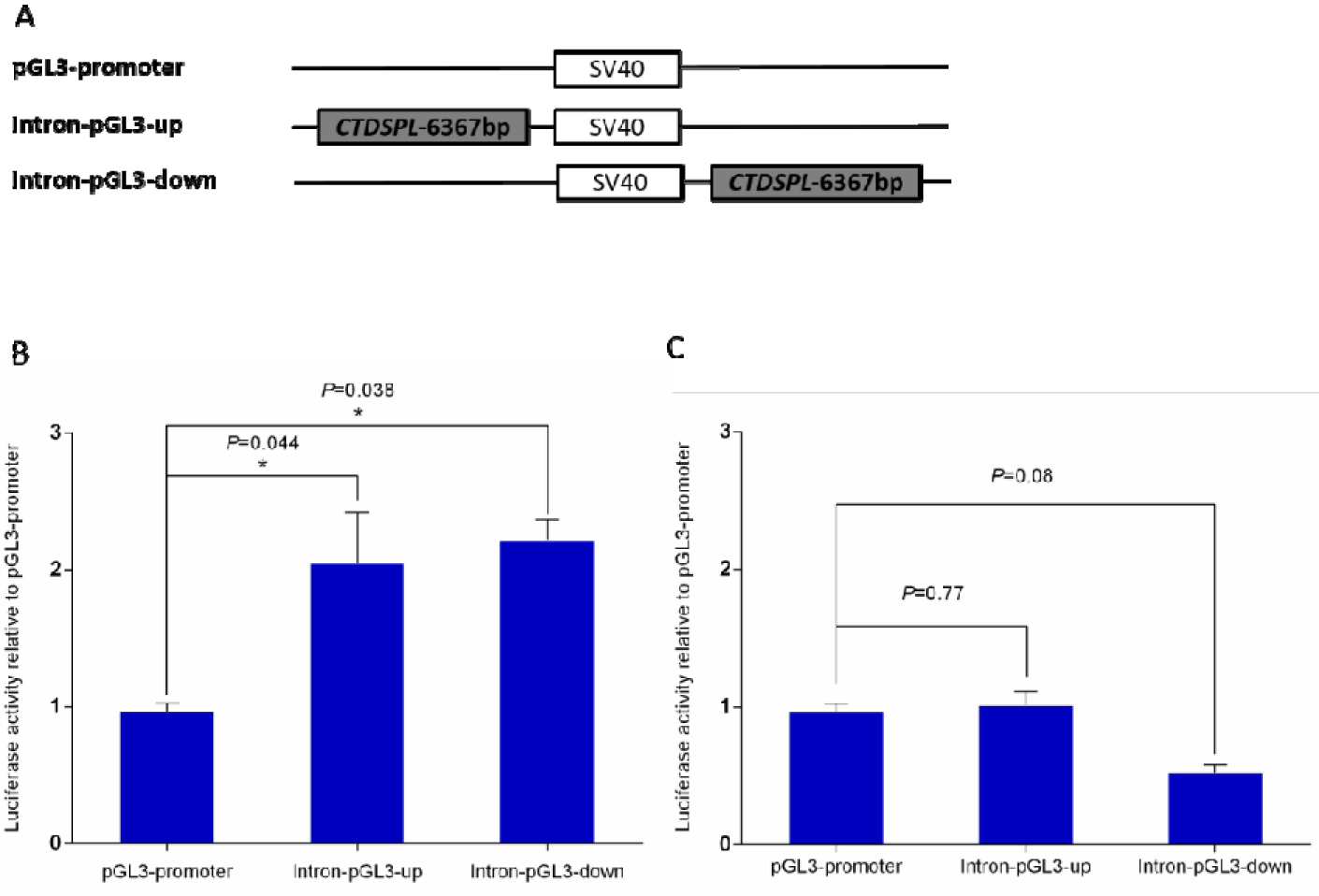
(A). The structure of pGL3 constructs of Luciferase assay. (B)Luciferase signals of three pGL3 constructs in HEK293T cells which overexpressed ZNF263.(C)Luciferase signals of three pGL3 constructs in HEK293T cells which did not express ZNF263. Statistical significance between experimental groups was assessed using a T-test.

**Figure 5.**
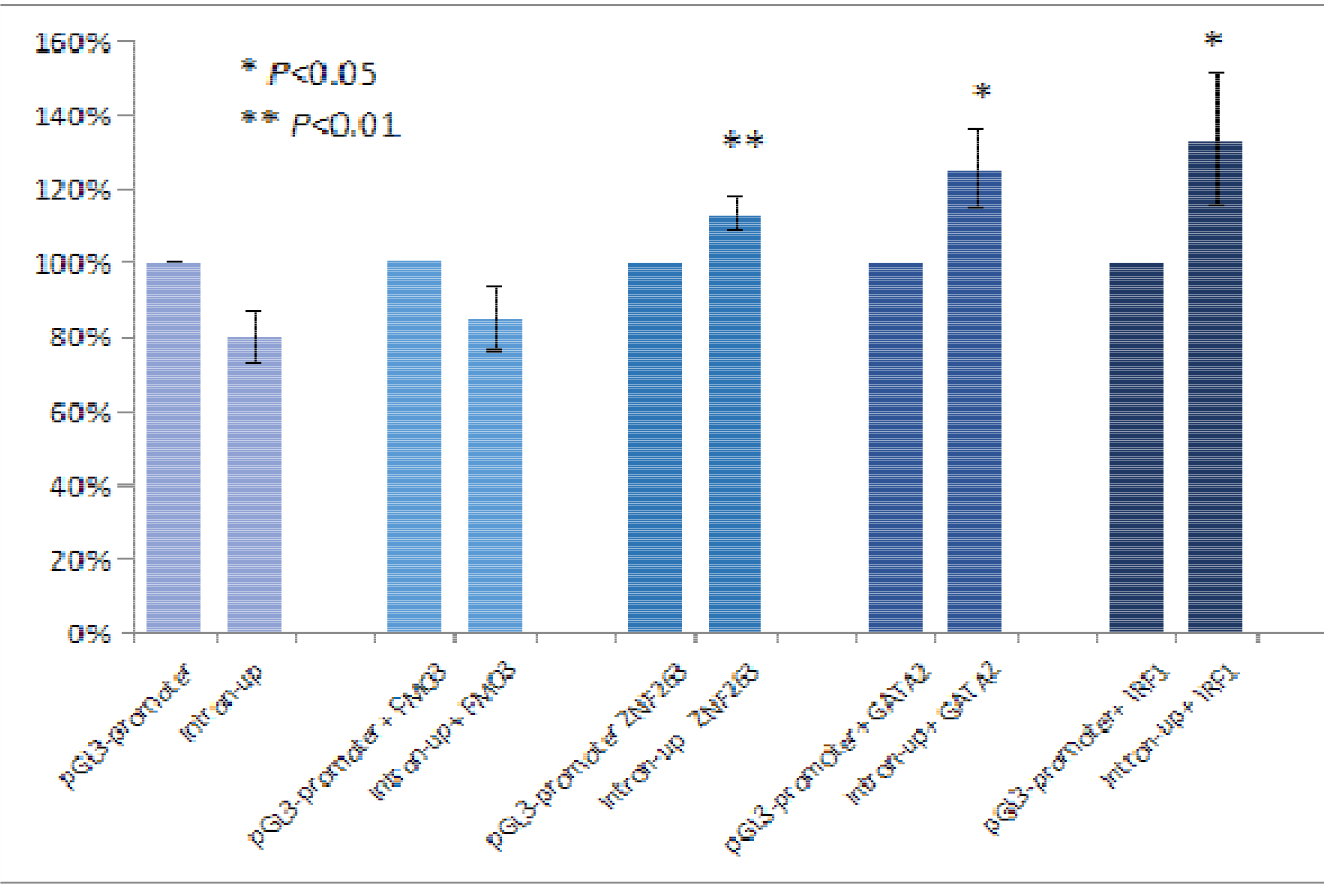
Luciferase signals of two pGL3 constructs in HEK293T cells which overexpressed FMO3, ZNF263, GATA2 and IRF1, respectively. Statistical significance between experimental groups was assessed using a T-test.

### Function of *CTDSPL* in cervical cancer

To determine the function of *CTDSPL* in human cervical cancers, we ectopically expressed *CTDSPL* in HeLa cells lacking *CTDSPL* (**Methods**). HeLa cells expressing *CTDSPL* showed a significant decrease in colony-forming ability compared with cells expressing vector control (**Figure 6A, B**). Next, we examined the role of *CTDSPL* on tumour growth (**Methods**). HeLa cells expressing *CTDSPL*, or vector control, were injected subcutaneously into immunocompromised mice. Compared with control groups, mice injected with HeLa cells expressing *CTDSPL* exhibited a significant reduction in tumour volume (**Figure 6C-E**). Histologic analysis confirmed that all mice bearing tumours expressing *CTDSPL* showed a significant down-regulation of Ki-67 (**Figure 6F**). Moreover, we used two distinct short hairpin RNAs (shRNA1 and shRNA2; hereafter referred to collectively as shCTDSPL) to ablate *CTDSPL* expression in luciferase-expressing End1/E6E7 cells (**Figure 6G**). We then examined the effect of *CTDSPL* depletion on tumourigenicity by subcutaneous injection into mice with End1/E6E7 cells transduced with shCtrl or shCTDSPL (**Methods**). We observed that the mice implanted with control cells were viable and did not develop tumours when sacrificed, whereas *CTDSPL*-depleted immortalized End1/E6E7 could form tumours in NOD-SCID mice (**Figure 6H**). The tumor burden in the whole cohort was shown in Figure 3I.

**Figure 6.**
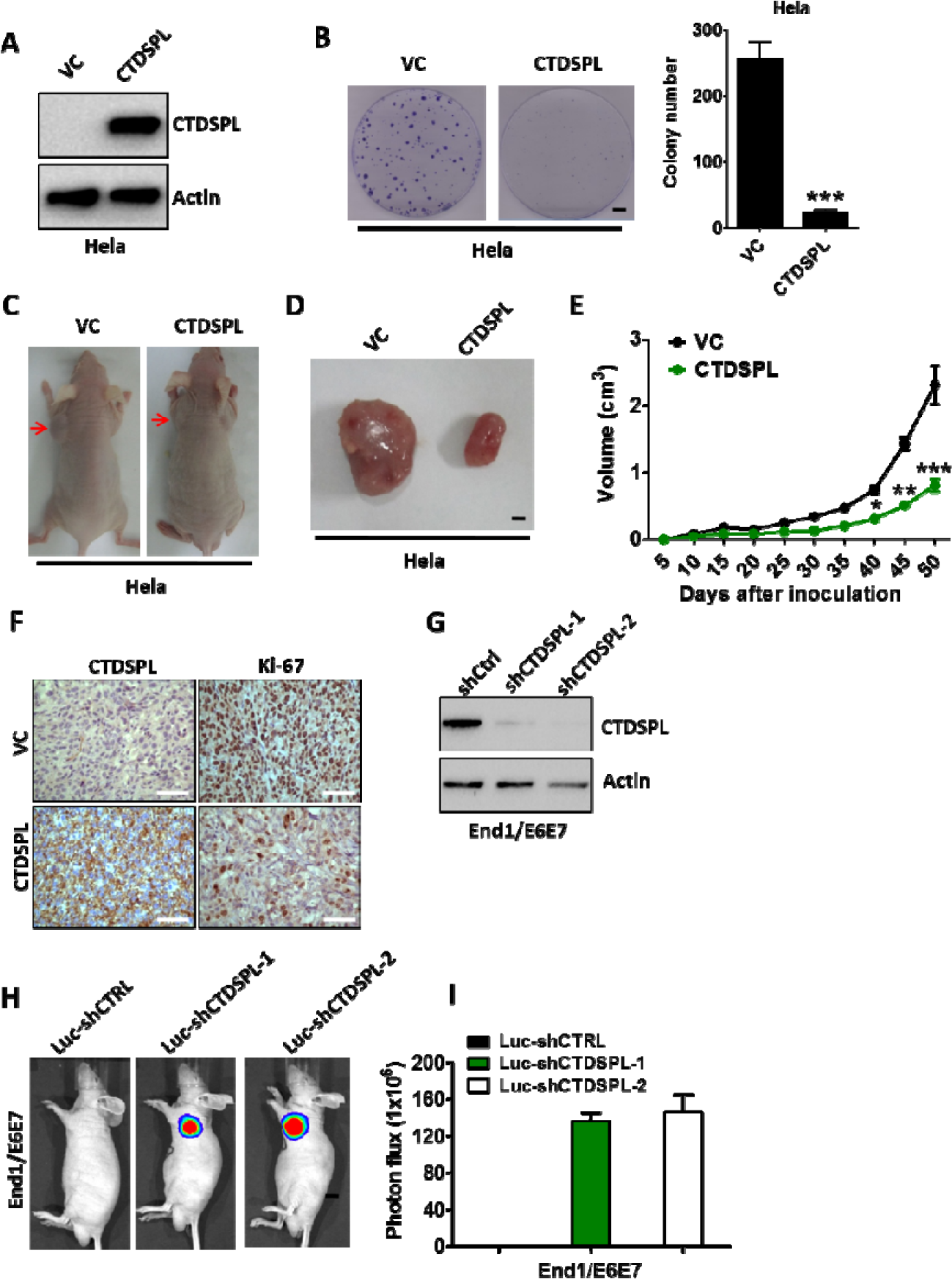
**(A)**. Western blot analysis of *CTDSPL* expression in Hela cells transfected with pcDNA3.1-CTDSPL or vector control, respectively. Actin served as the loading control. **(B)**. Colony formation ability of Hela cells after transfection of pcDNA3.1-CTDSPL or vector control. Data are means of three independent experiments ± SEM. ***, *P*< 0.001. **(C)** and **(D)**. Representative images of xenografts and tumours originated from Hela CTDSPL-overexpressing or vector control cells on the 50 days are shown. **(E)**. Growth curve of CTDSPL-overexpressing or vector control cells-derived subcutaneous tumour xenografts. *, *P* < 0.05; **, *P*< 0.01; ***, *P* < 0.001. *(F)*. The level of *CTDSPL* and Ki-67 was examined by IHC analysis in CTDSPL-overexpressing or vector control cells-derived tumour xenografts. Representative images of IHC are shown. **(G)**. Western blot analysis of *CTDSPL* expression in End1/E6E7 cells transfected with two independent luciferase-encoding *CTDSPL* shRNA or control shRNA. Actin served as the loading control. **(H)**. Representative pseudocolour bioluminescence images of mice bearing CTDSPL-depleted End1/E6E7 cells or control cells. **(I)**. The tumor burden in the whole cohort of 8 mice/group. Statistical significance between experimental groups was assessed using a T-test.

## Discussion

We have performed the first genome-wide CNV study of cervical cancer. We found that a 6367bp deletion in intron 1 of *CTDSPL* was associated with 2.54-fold increased risk of cervical cancer. This fragment is likely to enhance *CTDSPL* transcription with presence of transcriptional factor ZNF263, GATA2 or IRF1. *In vitro* and *in vivo* studies indicate that *CTDSPL* is a new tumour suppressor gene for cervical cancer.

No frequency data can be found for the detected 6367bp deletion in intron 1 of *CTDSPL* in the 1000 Genome Project (Genomes Project et al. 2010). However, the CNV esv2676043 (chr3:37978417-37986927) identified in the 1000 Genome Project which covers the *CTDSPL*-6367bp deletion showed frequency of 0.02, 0.06, 0.09 and 0.05 in the African population, American population, European population and Asian population, respectively.

*CTDSPL* encodes a protein phosphatase that dephosphorylates RB1, halting the cell cycle at the G1/S boundary, thereby controlling cell proliferation (Luo et al. 2015). It is conserved from yeast to human. *CTDSPL* is frequently deleted in cervical tumour and cervical intraepithelial neoplasia (CIN). The deletions in *CTDSPL* have been found to be significantly higher in squamous cell carcinoma (SCC) with metastases than in SCC without metastases, and decreased expression was more frequent in SCC with metastases as compared to SCC without metastases (Kashuba et al. 2009). Reduced expression of *CTDSPL* in SiHa and CaSki along with tumour suppressive ability has been reported in both *in vitro* and *in vivo* systems (Anedchenko et al. 2007). The promoter of *CTDSPL* is highly methylated in cervical cancer (Mitra et al. 2010b). Furthermore, *CTDSPL* deletion is associated with poor prognosis of cervical cancer patients (Mitra et al. 2010b).

Up to now, little is known about the function of ZNF263, except that it was predicted to have a repressive effect on gene transcription and often binds intragenic regions (Frietze et al. 2010). Our finding that ZNF263 upregulated *CTDSPL* provides new clues to ZNF263 function. Chen et al. reported that ZNF263 was upregulated in the blood of hepatocellular carcinoma (HCC) patient compared with the healthy volunteers (Chen et al. 2017), which implies that ZNF263 may play a role in the pathogenesis of HCC. GATA2, a member of GATA family of transcription factors, is expressed principally in hematopoietic cell lineages, with a particularly prominent expression in early progenitors, as well as in megakaryocytes and in mast cell lineages (Vicente et al. 2012). Mutations in this gene have been related to various hematological malignancies, particularly in MDS/AML (myelodysplastic syndrome/acute myeloid leukemia) familial aggregations (Kazenwadel et al. 2012). Recently, the functions of *GA TA2* as an oncogene in different types of human cancer have also been reported (Kumar et al. 2012; Wang et al. 2012; Chiang et al. 2014). For example, *GATA2* silencing could decrease cell migration and tissue invasion in prostate cancer (22021428Bohm et al. 2009). Willman et al. presented convincing evidence that *IRF1* was tumor suppressor gene (Willman et al. 1993). Loss of heterozygosity (LOH) at the IRF1 locus occurs frequently in human gastric cancer (Tamura et al. 1996). In addition, Harada et al. observed alternative splicing of *IRF1* mRNA, producing nonfunctional IRF1 protein at high frequencies in patients with myelodysplastic syndrome and acute myelogenous leukemia (Harada et al. 1994).

Several limitations should be addressed. First, the technical validation of CNVs in the present study was not sufficient. A random sample of both cases and controls should have been taken for technical validation, oversampling those with the deletion. Unfortunately, we didn’t have access to the DNA samples of the controls. Therefore, we could only oversample subjects with the deletion and randomly selected subjects without the deletion for technical validation. Second, there are no SNPs on the Illumina HumanOmniExpress BeadChip that cover *DEFB4* in our study. Therefore, we could not evaluate the association between *DEFB4* copy number and susceptibility to cervical cancer.

In summary, we found that a 6367bp deletion in intron 1 of *CTDSPL* was associated with 2.54-fold increased risk of cervical cancer. This CNV is one of the strongest genetic risk variants identified so far for cervical cancer. This deletion removes the binding sites of ZNF263, GATA2 and IRF1, and hence downregulates the transcription of *CTDSPL. In vitro* and *in vivo* studies suggest that *CTDSPL* is a tumour suppressor gene for cervical cancer.

## Methods

### Study population

Subjects included in the discovery phase were from a GWAS of cervical cancer in the Swedish population. The details of population and quality controls have been described elsewhere (Chen et al. 2013). Briefly, subjects included in the discovery phase were from two studies, the CervixCan I study and the TwinGene study. CervixCan study included two parts, i.e. CervixCan I study that comprised cases who are the sole participants of their family and CervixCan II study that comprised individuals with more than one first-degree relative also participating. 766 sole participants (720 CIS and 46 invasive carcinoma) from the CervixCan study were included in the discovery phase. The TwinGene study was a population-based Swedish study of twins born between 1911 and 1958. In total, 9,896 subjects were genotyped consecutively with those from the CervixCan I study using Illumina HumanOmniExpress BeadChip (731,422 SNPs) at the SNP&SEQ Technology Platform Uppsala, Sweden. Among these subjects, 309 unrelated cervical cancer cases (288 CIS and 21 invasive carcinoma) were further included in the discovery phase. One female singleton was then randomly selected from each twin pair without cervical cancer, resulting in 4,014 unrelated cervical cancer-free females who were included as controls. After stringent quality control (Chen et al. 2013), there were genotyping data in the discovery phase including 632,668 SNPs with an overall call rate of 99.92 % in 1,034 cervical cancer patients (971 with carcinoma *in situ* (CIS) and 63 with invasive carcinoma) and 3,948 control subjects. The replication series comprised 1,396 cervical cancer patients (1,265 CIS and 131 invasive carcinoma) from CervixCan II study and 1,057 controls. All the subjects were of Swedish descent. Informed consent was obtained from all subjects and each study was approved by the regional ethical review board in Uppsala, Sweden.

### CNV detection

CNV coordinates were identified using both Penncnv and Quantisnp (Colella et al. 2007; Wang et al. 2007). Both of the two algorithms are based on a Hidden Markov Model (HMM), using intensity files generated by GenomeStudio software from Illumina. QuantiSNP2.0 is based on an objective Bayes HMM and takes into consideration log R Ratio (LRR) as well as B allele frequency (BAF) of each SNP. The PennCNV algorithm incorporates additional information including the population frequency of the B allele (PFB) and the distance between adjacent SNPs. To reduce false positive calls due to genomic waves, GC-content adjustment was performed to correct for the bias in both analyses (Diskin et al. 2008). The default setting was used for both algorithms.

### Quality control

The initial sample quality control has been described elsewhere (Chen et al. 2013). All the SNPs that pass quality control were included in CNV calling. The proportion of the array that was informative for the CNV calling was 632,668/731,422=86.5%. Samples were further filtered based on three additional criteria: individuals with more than 40 CNVs; an absolute value of GC wave factor (GCWF) larger than 0.02 or a standard deviation of LRR > 0.30, as recommended by PennCNV; a genome-wide LRR SD obtained from QuantiSNP greater than 3.50. To select CNVs with high confidence for downstream analyses, the following criteria were applied: (i) with a maximum Bayes factor > 10 predicted by Quantisnp, (ii) called by both Quantisnp and PennCNV, and the breakpoints identified by the two algorithms should be within 2bp difference, (iii) with a physical length greater than 1kb and spanning three or more contiguous probes. As to every specific gene/region that we were interested, more profound quality control was performed. Samples with inconsistent CNV calls from two algorithms were further removed. The potential for population stratification was investigated by PCA undertaken with the EIGENSTRAT package (Allen-Brady et al. 2006). 9 significant eigenvectors were identified based on the Tracy-Widom statistic (*P*<0.05) but none of them was significantly associated with case-control status (All *P*>0.05), suggesting that population stratification is not a confounder in our study.

### Global burden analysis

We assessed the genome-wide CNV burden in patients and controls based on the number of CNVs (all CNVs, all genic CNVs, rare CNVs and rare genic CNVs) per genome. CNVs with a frequency **≤** 1.0% in our dataset were defined as rare CNVs. Genic CNVs were defined as CNVs overlapped with one or more genes (Genes were determined by RefSeq annotations (UCSC, Feb 2009, GRCh37/hg19) and gene boundaries were extended with a 10 kb flanking region on either side (Pinto et al. 2010)). We first evaluated the burden of deletions and duplications together, followed by one or the other separately. Significant differences were determined by 1 × 10^6^ permutations via PLINK (Purcell et al. 2007).

### Gene-based analysis

The numbers of patients and controls in whom a given gene was affected by CNVs were counted and compared. Permutation tests (1 × 10^6^ iterations) were carried out for individual genes by PLINK and permutation correction for multiple testing for all genes was performed with the max (T) permutation (mperm) option of PLINK (Purcell et al. 2007). In addition, odds ratio (OR), 95 % confidence interval (CI) and *P* values for specific CNV-disrupted genes for the risk of cervical cancer were calculated by logistic regression in the allelic test using the SAS 9.3 software. Bayes false discovery probability (BFDP) calculation was performed to reduce the probability of false-positive findings from GWAS stage. Two levels of prior probability (0.05 and 0.005) was selected and prior OR of 1.5 was adopted, which was suggested by Wachoder et al for a detrimental variant. BFDP <0.5 was used to indicate noteworthy findings (Wakefield 2007).

### Technical Replication

CNVs in *CTDSPL* and *NEDD4L* were technically validated using SybrGreen real-time quantitative PCR (RT-qPCR) on the ABI7900HT Fast Real-Time PCR System (Life Technologies, CA, USA). The Ct values were obtained with the SDS software v2.3 (Life Technologies). Copy numbers were calculated by the ΔΔCt method with normalization against both the reference gene and the samples without the corresponding CNV. In addition, ATPase 13A4 (*ATP13A4*) was excluded from the study as its CNV segment contained highly repeated sequences, which hindered successful design of efficient and specific primers. The following primers were used: *CTDSPL* (forward 5’-CTGGTGCTTTGAAGATACGG-3’, reverse 5’-AGCAATAGGCTTACAGAGGG-3’), *NEDD4L* (forward 5’-TGCTACTGAC AGCCTAAATC-3’, reverse 5’-GGACCTCTGAGCCATAAAAG-3’) and *RNase P* (reference gene with two copies in diploid human genome, forward 5’-TATTCACAAAGAGCCCAGAG-3’, reverse 5’-GAAGGGTATGGGAAAACAAG-3’). The PCR reaction were performed in triplicates and comprised 10 ng of genomic DNA, 200 nM of each primer and PowerSYBR^®^ Green PCR Master Mix (Cat. No. 4367659, Life Technologies). Samples were denatured at 95°C for 10 min followed by 40 cycles of 15 sec denaturation at 95°C and 1 min annealing at 60°C.

### Independent Replication

*CTDSPL* and *NEDD4L* CNVs were independently determined in the replication cohort by using Custom Taqman Copy Number Assays (Life Technologies). Furthermore, CNVs of all the SybrGreen RT-qPCR validated samples were double validated and separately included as controls in the Taqman assays during each run. Data were obtained by the SDS software v2.3 (Life Technologies) and copy numbers were calculated using the Copy Caller v2.0 software with a calibrator sample without deletion of the CNV segments. The following oligonucleotides were used: *CTDSPL* forward primer 5’- GGTACAAATCTGATCCCGTCACT -3’, *CTDSPL* reverse primer 5’-CTTGCAAGCCATGGAGATGAG -3’, *CTDSPL* FAM-labeled probe 5’- CCCTTTTCCATAACATCAAATCC -3’, *NEDD4L* forward primer 5’-TTGGGTAAATCATGGCTTAAAACTCTCA -3’, *NEDD4L* reverse primer 5’-TCTGAATGCAGGGTGGGAAATAAAA -3’, *NEDD4L* FAM-labeled probe 5’-CTAGGTTCTTGCACATCTTTGC -3’. The *RNase P* primers and VIC-labelled probe included in the Taqman Copy Number Reference Assay (Cat. No.4403328, Life Technologies) were used as references and mixed with either the *CTDSPL* primer-probe mix or the *NEDD4L* primer-probe mix. Reactions were performed in duplicates using 10 ng of genomic DNA, target and reference gene primer-probe mix and Taqman Universal Master Mix (Cat. No. 4326614, Life Technologies) in a duoplex format, by using the above-mentioned amplification cycles on the ABI7900HT Fast Real-Time PCR System.

### Replication and combined statistical analysis

To account for the correlation between related subjects in the replication and combined datasets, the association analysis was performed using PedGenie that allows for a mixture of pedigree members and independent individuals to be analyzed together (Allen-Brady et al. 2006). A Monte Carlo approach is employed to generate an empirical null distribution from which significance for a statistic of interest can be determined. Empirical 95 % CIs and *P* values were calculated using 1×10^19^ simulations.

### Correlation between CNV and gene expression

293 CSCC and endocervical adenocarcinoma from TCGA (http://www.cancergenome.nih.gov) with both CNV data and RNA-seq data available were included in this study. Level 3 CNV data detected from Affymetrix SNP 6.0 microarrays were downloaded. Segment mean values of the segments covering the 6.4kb deletion were extracted according to chromosome positions, and the copy number was calculated as (copy number value = 2*2^segment mean values). Copy number value between 1.7 and 2.3 were defined as no copy number variation, whereas <1.7 was defined as deletion and >2.3 was defined as duplication. In addition, RSEM-normalized results for RNA-seq data were downloaded. The association between the CNV of *CTDSPL* and expression of host gene and surrounding genes within 200kb was calculated by one-way ANOVA, Spearman correlation and linear regression analyses.

### Cell culture

HEK293T was cultured in Dulbeccos modified Eagle’s medium (DMEM, Gibco) supplemented with 10 % fetal bovine serum (FBS, Gibco). Human epithelial cervical cancer cells (HeLa) purchased from America Type Culture Collection (ATCC) were cultured in 75 cm^2^ flasks in Dulbecco’s modified Eagle’s medium (DMEM) (Sigma-Aldrich) supplemented with 10 % (v/v) fetal bovine serum, 100 IU/mL penicillin and 100 mg/mL streptomycin. End1/E6E7 cells (human endocervical cells immortalized with human papillomavirus E6/E7) were obtained from ATCC and cultured in 75 cm^2^ flasks in keratinocyte serum-free medium supplemented with keratinocyte growth supplement, 100 U/ml penicillin and 100 mg/ml streptomycin. All cells were maintained at 37°C in a 5% CO_2_ incubator.

### Lentiviral plasmid construction, lentiviral production and over-expression of ZNF263 cell line screening

The gene *ZNF263* was PCR-amplified from a human cDNA library and was fused into a lentiviral vector containing an EF1α promoter, which co-expressed an EGFP fluorescence protein from a CMV promoter. The lentiviruses of ZNF263 were obtained by co-transfection with three viral packaging plasmids pLP1, pLP 2 and pLP VSV-G into HEK293T using CaCl2 transfection. Virus was harvested 48 hr post-transfection. For viral transduction, cells were incubated with culture-medium-diluted viral supernatant for 24 hr. At 72 hr following transduction, the EGFP-positive population was reached to more than 90 %.

### Plasmid construction and luciferase assays in HEK293T stably overexpressing ZNF263

The *CTDSPL* intron 1-6367 bp fragment was PCR-amplified from 293T genomic DNA. To explore directionality of the regulatory element, we cloned the fragment upstream and downstream of SV40 of the pGL3-promoter vector (Promega), respectively. Inserts in each construct were verified by sequencing. Primer sequences are available on request. Constructs were transfected with equimolar amounts (800 ng) for luciferase reporter plasmids into HEK293T and HEK293-ZNF263 using lipofectamine 2000 reagent (Invitrogen), according to manufacturer’s instructions. Cells were harvested after 48h. Luminescence activity was measured with a Berthold lumat LB9507. Assays were performed in triplicate wells. Data represent at least three independent experiments. Statistical significance between experimental groups was assessed using a T test analysis. All analyses were performed using SAS 9.3, and a two tailed *P* value <0.05 was considered significant.

### Plasmid construction and luciferase assays in HEK293T stably overexpressing ZNF263

The *CTDSPL* intron 1-6367 bp fragment was PCR-amplified from HEK293T genomic DNA. To explore directionality of the regulatory element, we cloned the fragment upstream and downstream of SV40 of the pGL3-promoter vector (Promega), respectively. Inserts in each construct were verified by sequencing. Primer sequences are available on request. Constructs were transfected with equimolar amounts (800 ng) for luciferase reporter plasmids into HEK293T and HEK293-ZNF263 using lipofectamine 2000 reagent (Invitrogen), according to manufacturer’s instructions. Cells were harvested after 48h. Luminescence activity was measured with a Berthold lumat LB9507. Assays were performed in triplicate wells. Data represent at least three independent experiments. Statistical significance between experimental groups was assessed using a T test analysis. All analyses were performed using SAS 9.3, and a two tailed *P* value <0.05 was considered significant.

### Plasmid construction and luciferase assays in HEK293T transiently overexpressing FMO3, GATA2, IRF1and ZNF263

*FMO3*, *GATA2* and *IRF1* cDNA were synthesised by Sangon Biotech (Shanghai, China) and were cloned into pcDNA3.1 vectors which contain a CMV promoter, respectively. *ZNF263* was PCR-amplified from a human cDNA library (Thermo Scientific CCSB-Broad Lentiviral Expression Library) and was fused into a pcDNA3.1 vector. The *CTDSPL* intron 1-6367bp fragment was PCR-amplified from HEK293T genomic DNA, and then was inserted upstream of the promoter -luc+ transcriptional unit of pGL3-promoter vector (Promega). All constructs were verified by sequencing. Primer sequences are available on request. Constructs were co-transfected with equal weight amounts (500 ng) of luciferase reporter plasmids and equal weight amounts (500 ng) of cDNA over-expression plasmids into HEK293T cells using lipofectamine 2000 reagent (Invitrogen), according to the manufacturer’s instructions. Luciferase expression was normalized to 50 ng Renilla luciferase expression (pRL-SV40). Cells were harvested after 48h. Luminescence activity was measured with a Berthold Centro LB 960 Microplate Luminometer. Assays were performed in fourfold wells. Data represent at least three independent experiments. Statistical significance between experimental groups was assessed using T test. All analyses were performed using SAS 9.3, and a two tailed *P* value <0.05 was considered significant.

### Plasmid construction and colony formation assay

*CTDSPL* cDNA fragment was amplified from Human Umbilical Vein Endothelial Cells (HUVEC) mRNA by RT-PCR. The product was extracted, purified and digested with Hind III and Xho I, and then was inserted to myc-tag fusion vector pcDNA3.1/myc-his-A. The base sequence of recombinant pcDNA3.1/myc-CTDSPL plasmid was in accordance with human *CTDSPL* cDNA fragment by agarose gel electrophoresis and DNA sequence analysis. *CTDSPL* shRNAs were purchased from Santa Cruz Biotechnology. All oligonucleotides and plasmids were transfected into cells using Lipofectamine 2000 Transfection Reagent (Invitrogen) according to the manufacturer’s instructions.

Before genetic manipulation, cells were engineered to stably express firefly luciferase by transfection with pNifty-CMV-luciferase and selection with 500 μg/ml Zeocin. 1×10^3^ cells were independently plated onto 60-mm tissue culture plates. After 10-14 days, visible colonies were fixed with 100 % methanol and stained with 0.1 % crystal violet in 20 % methanol for 15 min. Colony-forming efficiency was calculated as the number of colonies/plated cells × 100 %.

### Immunohistochemistry

The immunohistochemistry assay was conducted on nude mouse xenograft tumour tissues to detect and score CTDSPL and Ki-67 expression using methods described previously (Luo et al. 2015). CTDSPL and Ki-67 antibodies was obtained from Novus Biologicals. Statistical significance between experimental groups was assessed using a T-test. All analyses were performed using SAS 9.3, and a two tailed *P* value < 0.05 was considered significant.

### Tumour xenograft models

All animal experiments were conducted with the approval of the Shanghai Jiao Tong University Institutional Committee for Animal Research and in conformity with national guidelines for the care and use of laboratory animals. Cells infected with a luciferase-encoding lentivirus (1 × 10^6^) were inoculated subcutaneously into nude mice. There were 8 mice in each experimental group. Tumour volume (V) was monitored by measuring tumour length (L) and width (W) with callipers and then calculated with the formula (L×W2) × 0.5. Statistical significance between experimental groups was assessed using a T-test. All analyses were performed using SAS 9.3, and a two tailed *P* value <0.05 was considered significant.

## Acknowledgements

The authors thank all of the participants in this research and support staff who made this study possible. We are grateful to all pathology clinics that enabled access to their archives. Illumina genotyping was performed by the SNP&SEQ Technology Platform in Uppsala. The platform is part of Science for Life Laboratory at Uppsala University and supported as a national infrastructure by the Swedish Research Council.

